# Aviti Sequencing and Marker Gene Data Analysis

**DOI:** 10.64898/2026.02.06.704475

**Authors:** Trevor J Gould, Madeline Taylor, Cara Santelli

**Affiliations:** Minnesota Supercomputing Institute, University of Minnesota, Minneapolis, MN, USA; Department of Earth and Environmental Sciences, University of Minnesota, Minneapolis, MN, USA; BioTechnology Institute (BTI), University of Minnesota, Minneapolis, MN, USA

**Keywords:** Aviti, 16S, Marker Genes, Sequence Data, Chimeric Reads, Clustering

## Abstract

Accurate identification of microbial species in complex populations and communities relies on the isolation of representative marker 16S, ITS, and 18S sequences through the use of DNA extraction, PCR, and sequencing. Aviti sequencing has brought an improvement in the read quality and depth of marker gene sequencing technology. Quality scores exceeding Q40 representing highly accurate sequencing allows researchers to ask more questions of their marker gene data. However, this improvement in quality and throughput also brings with it a surprising increase in diversity of amplicon sequencing variants (ASVs) making further analysis and comparisons to previous studies on Illumina platforms challenging. This increased diversity causes downstream processing issues, including an over-reporting of chimeric ASVs. Here we identify this problem and put forward straightforward solutions to retain counts and reduce technically introduced diversity, as well as tying chimeric read identification to minimum parent distance. Through the use of synthetic mock samples, we discovered that erroneous ASVs are systematically substitution errors introduced by the upstream PCR methods. This error can be reduced significantly bioinformatically through clustering of ASVs within 99% similarity. Further we highlight technically introduced variation as a result of variable region length, sample misassignment, and sample biomass. Collectively, these results improve the similarity of Aviti and Illumina datasets for better comparisons of microbial studies from different platforms.

## Introduction

Sequencing technologies have gone through another generation of development with the advent of rolling circle amplification and the Avidity methods (Arslan et al.). Aviti sequencing has come into the market producing a large amount of higher quality short reads than has been seen from past generations of high throughput sequencing technology. With this improved quality and high number of total reads, issues specific to short read marker genes, such as chimeric detection and pipelines not designed for >Q40 reads, need to be overcome in order to process and analyze the data bioinformatically. Early adopters of the technology have noted the remarkably high number of amplicon sequencing variants (ASVs) in comparison to Illumina output (Benjjneb). This study investigates the source of this increased variability and points to several possible solutions to the problem.

In 2024, Element Biosciences released the Aviti sequencing platform, which produces an average phred-scaled quality score greater than Q40 (99.99% base calling accuracy) and produces as many as 1.5 billion reads of 2 × 300 per run (Arslan et al.). Additionally, the Aviti system scales quickly by using multiple lanes per machine at low cost. As a result, marker gene sequencing projects are only going to get bigger in terms of total reads. This scale creep results in a large increase in run times for processing of a commensurate amount of illumina data, due in no small part to an increase in ASV variation noted by early adoption researchers (Benjjneb). Quality improvements have several consequences; first, more sequences are retained so counts are higher, and second, when quality is so high, error-correcting-pipelines based on the presence of errors can run into unexpected issues.

Cloudbreak developed by Element Biosciences allows for the use of third party adapters in the use of Aviti sequencing (“Cloudbreak™ Is Here”). This increases the applications that can be used with the sequencing platform. Nextera adapters and Kapa-hifi-polymerase are commonly used in sample prep for PCR and illumina sequencing which can now be used in Aviti as well (Potapov and Ong). This improves the ability to compare sequencing methods between Illumina and Aviti, as ultimately the goal for most researchers is to be able to use Aviti as their primary platform and thus being able to compare to previous sequencing runs is essential.

Historically as the volume of illumina sequence data increased with technological improvements bioinformaticians faced a crisis of scale. While deeper coverage theoretically improved the ability to resolve rare taxa, it also produced more technical artifacts and at the time the scale of the data made computational resources a bottleneck in differentiating the signal from the technical noise (Pérez-Cobas). Chimeric sequence artifacts, hybrid sequences formed from amplicons priming off a different template, often appear as unique ASVs that need to be detected bioinformatically (de la Cuesta-Zuluaga, J., & Escobar, J. S.). Index-hopping, sequences miss-assigned to the wrong sample, frequently occurred in densely packed sequencing runs. In addition to methodological improvements, like unique dual indexing (Flörl et al.), bioinformatic pipelines used the correlation of quality scores and sequence abundance, to remove such technical noise (Callahan). As Aviti quality scores eclipse Q40 this quality to abundance relationship is challenged and researchers need to reassess the identification of chimeric reads and other technical noise.

In this study we used synthetic mock community samples included in Aviti sequencing runs with known reference sequences that contained a synthetic nine base pair insert, which allowed us to investigate potential sources of the increase in ASV variation. Because of the increase in sequence quality of Aviti, we are able to point to specific base pair substitution error patterns in the mock sequences and develop a method for remediating PCR induced error patterns. Finally, we compared large Aviti runs to prior Illumina runs of comparable sample sites and applied methodological improvements to increase the comparability of the two sequence data outputs. We hypothesize that clustering will be a beneficial step prior to the chimeric sequence removal step for Aviti data to group mutations with parent sequences and then remove chimeric sequences.

## Methods

### Full Datasets

### Sequencing

Two of the datasets (Miseq_1, Miseq_2) were generated on Illumina MiSeq instruments. For both of these datasets, amplicons were generated containing Nextera adapter regions following the process published by Gohl, et al. 2016. In summary, on-target template concentrations were determined using qPCR with the primers V4_515F_Nextera and V4_806R_Nextera, sample concentrations were normalized with molecular-grade water, and amplicons were generated by PCR. Adapters and unique dual indices were added with a second 10-cycle round of PCR and sample concentrations were normalized using SequalPrep kits (Thermo Fisher). Normalized samples were pooled into libraries, cleaned with magnetic beads, and diluted to ∼2 nM concentrations. Normalized libraries were loaded onto MiSeq instruments using V3 2×300 flow cells at 8 pM + 5% PhiX for sequencing.

Four of the datasets (Aviti_1:4) were generated on Element AVITI instruments. For all of these datasets, amplicons were generated identically as for amplicons sequenced on the Illumina MiSeq. Normalized libraries were loaded onto Aviti 2×300 Medium runs at 7 pM + 10% PhiX.

Each Aviti sequencing project run at the University of Minnesota Genomics Center includes a synthetic mock sample with the run as a control. For this study we have 21 samples from 21 different projects: 11 samples of V4 variable region, 3 of V3V4 variable region, 4 of V3V5 variable region, 3 of V5V6. The average read count is 397,408 sequences. These samples do not include the full sequencing runs however, just the synthetic mock samples.

### Sequence processing

Aviti sequence data was processed for adapters using Trimmomatic (Bolger et al.), followed by primer removal with cutadapt (Martin), a second round of adapter removal was performed using Trimmomatic as sequences with as many as five copies of a nextera adapters were present in the datasets. Following basic QC reads were processed with DADA2 (Callahan et al.) in R for further QC, ASV table creation, paired read merging, chimera removal, taxonomic estimation using current silva database for 16S or Unite for ITS datasets (Yilmaz et al.; Quast et al. 25; Glöckner et al.; Abarenkov et al.). Illumina datasets used the same methods to ease comparisons. Subsequent analysis was done in R (R Core Team) and unix bash command line utilizing custom code available on Github (see data availability).

### Synthetic Mock Samples

Mock sequence samples for Aviti projects were provided by the University of Minnesota Genomics Center (UMGC) and are included in all marker gene sequencing projects for ease of analysis (*UMGC* | *University of Minnesota Genomics Center*). The sequences contain 3 sets of 3 inserted bases to differentiate bacteria in the sample with that found in the study samples and provides an opportunity for mutation analysis of known sequences. Mutation detection analysis was done in R using the Biostrings package (Pages et al.). Stringdist function was utilized to compare the levenshtein distance of all sequences in the synthetic mock sequence with the known reference sequences. The pairwiseAlignment function was used to compare reference and subject ASVs for mutations on any sequence with two or less distance using a global alignment method, gapOpening = 5, gapextension =2. Detected mutations were tallied and all substitutions were included in table3 for the four full Aviti studies.

ASVs in the synthetic mock samples with a distance greater than zero and less than 5 to a reference were considered to be more likely to be from a reference sequence than to be a real biological sequence; with random mutations/PCR/sequencing error that made them increasingly similar to the mock insert sequence ASVs. These less-than-5 distance sequences are referred to here as erroneous ASVs for the purposes of the analysis. Sequence count of erroneous ASVs in the synthetic mock samples were noted and compared to the variable region that was sequenced.

### Clustering method

Clustering of ASV counts by sequence similarity merges counts of ASVs if the two sequences are within a set number of errors of each other. This retains the total sequence count and retains the ASV that has a higher initial count and removes the ASV, but not the sequence count, of the erroneous ASV. This method uses pwalign’s (Aboyoun P, & Gentleman) nucleotideSubstitutionMatrix set to match = 1 and mismatch = -3 as the substitution matrix for pwalign’s pairwiseAlignment function. This function was used with the options global alignment, gapOpening = 5, gapExtension = 2. Levenshtein distance, as opposed to hamming distance, is used here to allow for inserts and deletions causing different lengths of ASVs. All ASVs closer in distance than the minimum allowed errors were included in the clustered ASV. ASVs that clustered with multiple other ASVs were only merged with the ASV with the largest count. In this way, we were able to cluster out any ASV that was the result of a set number or less errors within the base call or PCR steps of the sequencing pipeline whilst keeping the sequence count data. It should be emphasized that one vs all pairwise sequence alignment of large sequence datasets is a significant computational task that becomes increasingly computationally expensive as the number of unique ASVs increases. This method of clustering will need bioinformatic improvements to run on datasets with more than an estimated 250,000 unique ASVs. We are exploring multi-stage pairwise comparisons to improve performance but that is beyond the scope of this work.

## Results and Discussion

The most readily observed issue to date with marker gene sequencing on the Aviti platform is the order of magnitude increase in ASV variation over Illumina. (Figure 1) shows the scale of the issue comparing unique ASVs from a pair of Illumina and Aviti Projects. Aviti results in orders of magnitude more unique ASVs than Illumina using the same sample. It is important to note that Aviti is also producing significantly larger numbers of sequences than Illumina in these projects. Ideally a comparison of Novaseq and Aviti would be done on the same samples here instead but the cost to produce such experiments is difficult to justify. There are however multiple comparisons of Miseq and Novaseq to indicate that while Novaseq does create additional diversity, the two methods are comparable, and certainly not an order of magnitude difference in diversity is observed (Han et al., Singer et al.). The majority of Illumina ASVs (91.1%) are shared with Aviti but, owing in part to the order of magnitude more ASVs in Aviti, conversely only a small percentage of ASVs in Aviti are shared with Illumina (3.7%). Rarefaction curves (Figure 2) reiterate this point with Illumina curves leveling off at an estimated species number of 3,000 while Aviti levels off at greater than 30,000 estimated species in Aviti_1. This makes direct comparisons challenging for alpha and beta diversity and increases the issues of technical sequencing artifacts. Table 3 further characterizes the overlap between Illumina and Aviti sequencing for both ASV variation and sequencing count. Encouragingly, a majority of the sequence count represented by ASVs in both methods are shared with the other; for example in table_2: 98.9% of Illumina reads and 68.3% of Aviti read counts are in shared ASVs. Clustering of unique ASVs at 99% identical and merging ASV counts of clustered ASVs could be used to maintain the total counts of the dataset and retain the more populous ASVs instead of obscuring with OTU designations. Alternatives to clustering, such as downsampling prior to sequence processing or rarefaction after sequencing processing, do not remove the technical variation in these datasets, but rather simply reduce its prevalence and use significantly less of the total counts.

**Figure 1.**
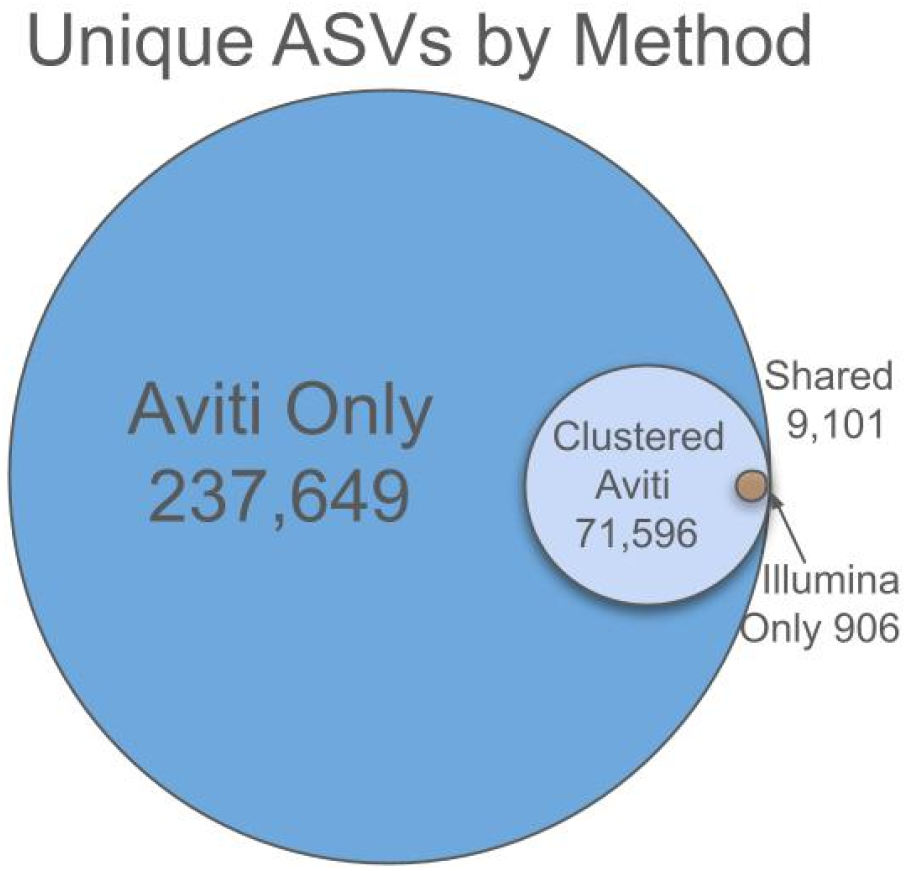
Unique ASVs in Aviti data are orders of magnitude larger than paired Illumina data. Shared ASVs between Aviti and Illumina are a larger proportion of total Illumina ASVs than Aviti. Clustering Aviti unique ASV data at 99% identity reduces proportional difference.

**Figure 2.**
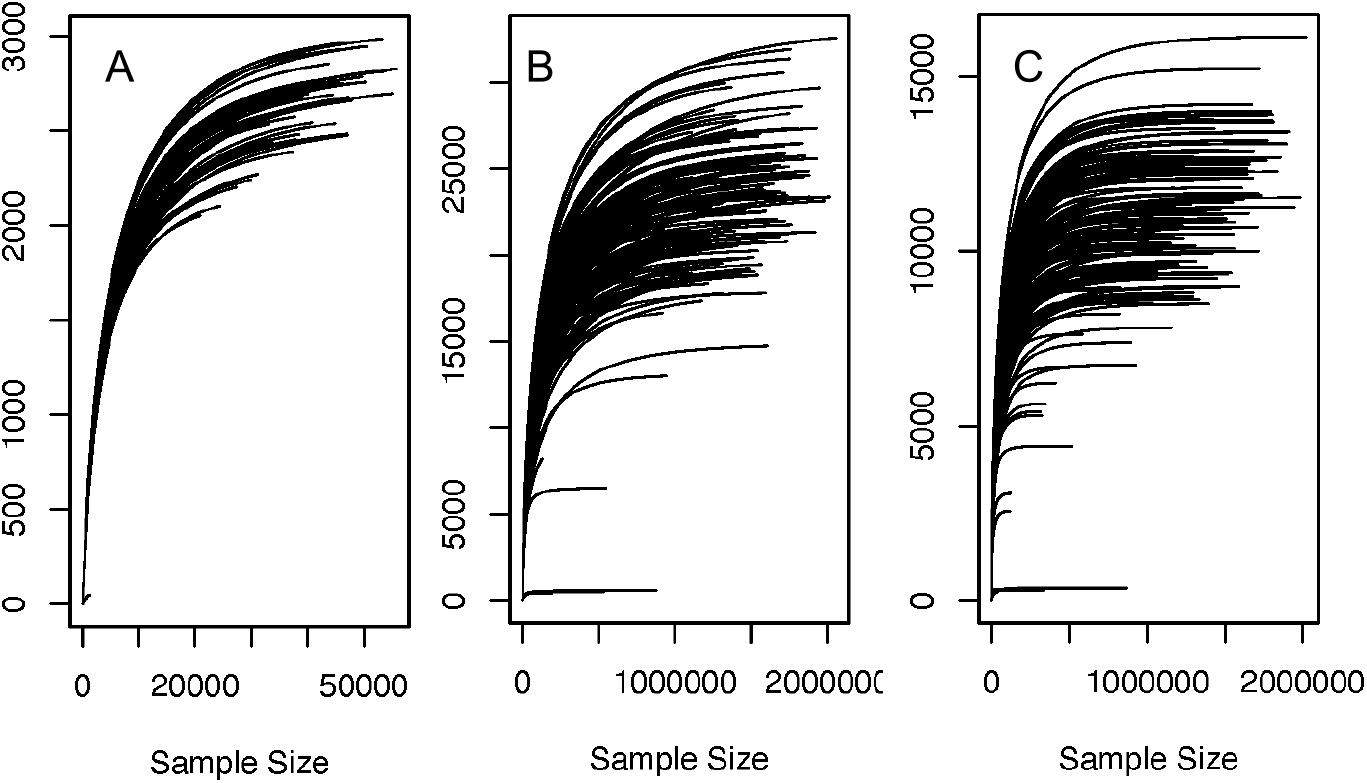
Rarefaction curves without replacement a) Illumina b) Aviti c) clustered Aviti There is an order of magnitude estimated species in Aviti vs Illumina data. Clustering aviti data reduces that difference but would also require additional filtering for Aviti data to be comparable to Illumina.

A second observation made about Aviti marker gene data is that in some, but not all, datasets, a large proportion of the unique sequences are removed at the chimeric sequence removal step (Benjjneb). Indeed we have observed the same thing; in one dataset 93% of the unique ASVs were removed at this step. Closer inspection of this Aviti dataset revealed that sequences removed in this way contain single base mutations from parent sequences. Comparisons of mutation rates vs chimeric sequence rates indicate that mutations occur more frequently (Lahr Katz). Chimeras are also more likely to have parents of more populous sequences indicating the randomness of the process (Lahr Katz). A second dataset with 31,997 unique ASVs had 70% ASVs removed (Table 3). Aviti synthetic mock samples contained 18 actual bacterial species and the same number of ASVs. In the four full datasets that were analyzed, the synthetic mock samples themselves contained an order of magnitude more unique ASVs. Pairwise alignment of all ASVs in the synthetic mock samples with reference sequences revealed a systematic bias in substitution errors in ASVs with two or less bases difference from reference sequences. Almost all of the substitutions were G to T and C to A in direction (Table 2). This is consistent with other findings in Illumina sequencing analysis of Nextera kapa hifi pcr error in high quality sequences (Shagin et al.).

**Table 1.** Full sequence datasets. Aviti_1 and Illumina_1 are from the same sample site, as are Aviti_2 and Illumina_2. Aviti_3 and Aviti_4 are included to improve the range of Aviti datatypes.

| ID | Samples | Total_Read_Pairs | Mean read-pairs per sample | PCR Target | Platform | Source |
| --- | --- | --- | --- | --- | --- | --- |
| Aviti_1 | 138 | 223,767,389 | 1,621,502 | Paired-end 300bp V4 | Aviti | Soil |
| Aviti_2 | 156 | 209,426,312 | 1,342,476 | Paired-end 300bp V4 | Aviti | Snow |
| Aviti_3 | 161 | 221,349,219 | 1,374,839 | Paired-end 300bp V4 | Aviti | Mouse |
| Aviti_4 | 90 | 106,198,562 | 1,179,984 | Paired-end 300bp V4 | Aviti | Human |
| Illumina_1 | 44 | 2,740,166 | 62,276 | Paired-end 301bp V4 | Miseq | Soil |
| Illumina_2 | 65 | 3,974,907 | 61,152 | Paired-end 301bp V4 | Miseq | Snow |

**Table 2.** Sequenced Aviti mock samples contain a majority of G-T and C-A substitutions.

| Substitution | Aviti 1 | Aviti 2 | Aviti 3 | Aviti 4 |
| --- | --- | --- | --- | --- |
| G-T | 456 | 73 | 185 | 99 |
| C-A | 213 | 28 | 100 | 59 |
| G-A | 58 | 2 | 2 | 0 |
| C-T | 24 | 0 | 2 | 0 |
| C-G | 4 | 2 | 0 | 0 |
| A-G | 3 | 0 | 0 | 0 |
| G-C | 2 | 1 | 0 | 0 |
| T-C | 1 | 1 | 0 | 0 |

**Table 3.** Results of clustering on paired Illumina and Aviti unique ASVs.

| Dataset | Unique ASVs before clustering |  |  | Unique ASVs after clustering |  |  |
| --- | --- | --- | --- | --- | --- | --- |
|  | Aviti | shared | Illumina | Aviti | shared | Illumina |
| Aviti_1\|Illumina_1 | 237649 | 9101 | 906 | 71596 | 6912 | 906 |
| Aviti_2\|Illumina_2 | 177516 | 13184 | 3822 | 116475 | 9235 | 4711 |
| Aviti_3 | 79,236 | NA | NA | 37,875 | NA | NA |
| Aviti_4 | 10839 | NA | NA | 5986 | NA | NA |

Synthetic mock samples from 21 other (not full runs) Aviti projects were also analyzed. The 21 16S synthetic mock samples were compared by variable region sequenced and it was found that the longer the variable region the higher proportion of the total count of the sample that was contained in erroneous ASVs. (Figure 3A). This pattern is consistent with the idea that PCR errors introduced in early cycles are more likely to accumulate in longer sequences, because the greater number of bases increases the probability of an error. Errors that occur early are subsequently amplified through later cycles, increasing the apparent abundance of those erroneous ASVs. We also observed that samples with higher total sequence counts in the synthetic mock communities contained more unique erroneous ASVs (Figure 3B). There is also a negative relationship between the total sequence count and the sequence count within the erroneous ASVs. (Figure 3C).

**Figure 3.**
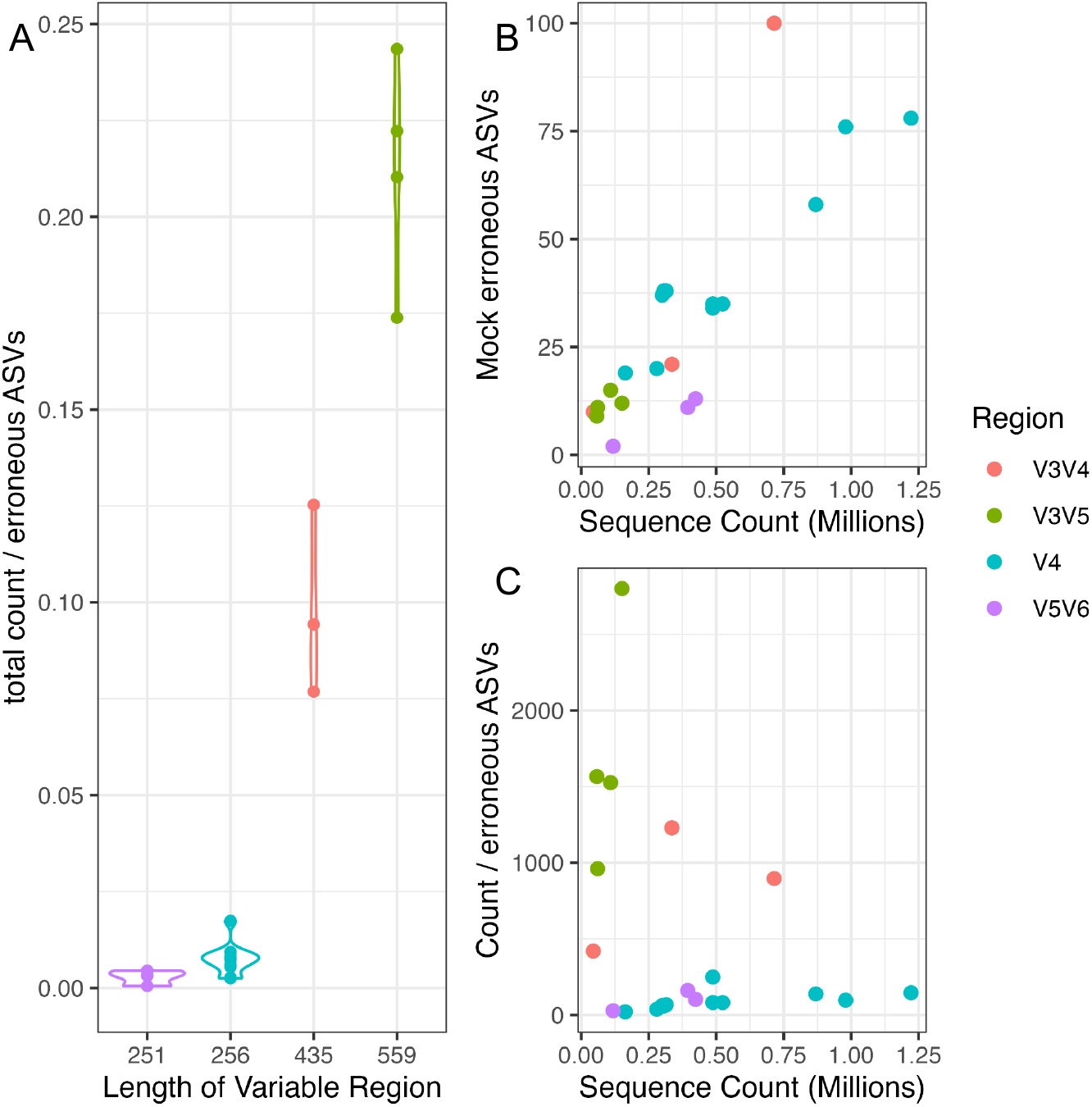
A) total count in each erroneous ASV increases by 16S variable region length. B) Number of Erroneous ASVs increases with sequence count. Indicating higher sequencing volume results in more errors. C) Negative relationship between total count and count per erroneous ASV. Lower volume sequencing has less total erroneous ASVs but those that do exist are higher count.

The method described here; clustering unique Aviti ASVs based on Levenshtein distance and merging sequence count data while retaining the ASV sequence of the higher count ASV; removes all ASVs in the synthetic mock sample within a set distance of the reference ASVs. Here we used a distance of less than 3 which is the closest approximation to a 99% clustering for a 253 base V4 ASV. In Aviti_1, we were able to retain all of the sequence count and reduce ASVs not found in the Illumina_1 run from 237,649 to 71,596 (Table3).

Again in Aviti_1 and Illumina_1, if just the ASVs present in both the Aviti and Illumina run are kept, this retains 68.3% and 98.9%, respectively, of the total count of sequencing, while 3.7% and 90.9% unique ASVs, respectively, are retained (Table 3). In Aviti_2 and Illumina_2, 77.5% of ASVs in the Illumina run were represented in the Aviti run. Conversely, shared ASVs represented 6.9% of the Aviti ASVs.Clustering increases Aviti ASVs shared to 8.8% and reduces the Aviti unique ASVs to 29% of its initial size (Table 3).

Clustering the Aviti and Illumina data from Aviti_2 and Illumina_2 combined 50,447 Aviti ASVs with those more abundant ASVs that were within two base pairs. 190,700 to 140,253: 68.3% of total count of sequencing in the Aviti run and 10,633 of the 17,006 or 62.5% ASVs from the illumina run are retained in the Aviti run. So clustering retains less Illumina ASVs in the Aviti run (77.5 to 62.5) but retains more Aviti sequences (6.9 to 7.6). 15% of illumina ASVs clustered with other illumina ASVs (2,551 of the 17,006) (Table 3). Of the 149 ASVs still present in the Aviti_2 synthetic mock sample after clustering, 19 are the mock sequences themselves, 72 are more abundant in real samples than the mock sample, indicating index bleeding or optical misidentification of the index sequence (Flörl et al).58 are more abundant in the mock sample than in the real samples; of which none had a distance of less than 17, some indicating additional primer/adapter contaminations, some indicating potential bacterial contamination (Davis et al, Ros-Freixedes et al). There is potential to use the synthetic mock samples as a cutoff for removing potentially contaminated ASVs. This would need to be explored on a case-by-case basis as to what threshold of counts is reasonable and expected for the study.

It is also notable that both sequencing methods, Illumina and Aviti, contain unique ASVs. There are two non-exclusive possible explanations; each method is able to capture some of the biological diversity present in the sample that the other method does not, or both methods are capable of introducing unique technical variation and error. Previous work in comparative methods of Illumina and 454 explored a similar dynamic finding that pseudo-beta diversity (between replicate differences) disappeared as sequencing counts increased (Smith et al, Hoisington et al). Given that both sequencing methods are using the same DNA extraction and PCR methods, and the vast majority of the method-specific ASVs are low count, the latter is a more likely case to explore in the future. Based on work presented here clustering large ASV datasets to 99% is preferable to setting arbitrary cutoffs for ASV removal. However, each dataset is uniquely dependent on the experimental design and processing choices. Further analysis of Aviti specific technical error is warranted.

### Misassignment

Another use for synthetic mock samples is the ability to detect potential index hopping, or sequencing reads that are assigned to the wrong sample (Flörl et al). Through the use of synthetic mock samples one can detect both synthetic sequences that were assigned to real samples and real sequences assigned to the synthetic sample, evidenced by a larger count of reads in real samples than in mock or blank samples. Estimated mis-assignment related to the synthetic mock sequences found in real samples is unchanged based on clustering and specific to the population environment (Table 4). This is useful to report as an estimate, and could be used as a threshold below which ASVs are considered suspect and zeroed; the mycology field uses this method for example with ITS sequencing in Illumina (Lindahl). It is not possible to determine from the data what caused the misassignments and careful future experiments need to be done to tease out the causes.

**Table 4.**
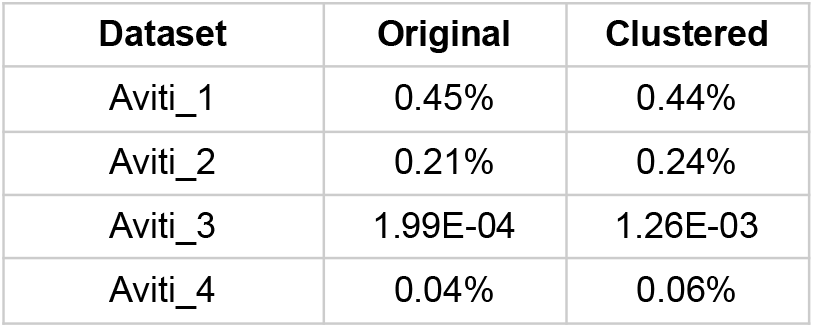
Percent of reads missassigned from synthetic mock samples to experimental samples before and after clustering in full Aviti datasets.

| Dataset | Original | Clustered |
| --- | --- | --- |
| Aviti_1 | 0.45% | 0.44% |
| Aviti_2 | 0.21% | 0.24% |
| Aviti_3 | 1.99E-04 | 1.26E-03 |
| Aviti_4 | 0.04% | 0.06% |

### Singletons

In Aviti_1, removing all ASVs with a count > 0 in only one sample keeps 96.8% of the sequence count and removes an additional 47,001 ASVs for a combined removal of 53.88% of the Aviti ASVs. There is a 60.6% correlation between the number of remaining singletons in a sample with the initial total unique ASVs in a sample (when controlling for the overlapping ASVs), but only a 19.5% correlation between the number of remaining singletons in a sample with the total count of the same sample. This indicates that singletons are not simply a product of sequencing volume. Indeed, clustering the same dataset to allow for 7 differences (97% clustering) in this dataset still has 29,996 singleton ASVs out of 69,596 total ASVs. However the shared ASVs represent 99.2% of the total sequencing; so functionally these ASVs are a minor proportion of the sequenced population, regardless of whether they are biological or just a technical byproduct of sequencing.

### Chimeric read detection

Aviti has an overabundance of error-induced reads with high quality that are treated as parents, which causes a majority of reads to be chimeric. This issue will be especially noticeable in a marker gene study where variation between bacterial species is low to begin with, and sequences are labeled chimeric even with single mutations from a parent as long as the full ASV matches any other pair of ASVs of a higher abundance. Therefore, reads from Aviti studies need to be considered as chimeric only when there are a higher number of differences between parent and potentially chimeric reads to avoid mischaracterizing reads. For this study, modifications to DADA2’s bimeric detection algorithm were made to allow setting a minimum difference between potential parent/child ASVs without enabling the parent one-off option. We recommend clustering at 99 percent before chimeric detection, then running a modified bimeric ASV detection that limits parent sequences by requiring parents to be a determined Hamming distance (the distance used in DADA2 chimeric detection) from child sequences. A similar method was developed with success by (Frøslev et al) called the LULU method. Interestingly, the LULU method also found limited benefit in removing singletons as compared to clustering. (Table 5) below is an example of a dataset where 70% of ASVs, and 10% of the total sequence count are removed by the chimera check using default dada2 settings. Determining the parent distance cutoff will require some customization based on the dataset being processed. Given that chimeric sequences are reported by Johnson et al. (2023) from 5 to 43% of indexes it’s difficult to give concrete advice in this space. In this example dataset, parent distances less than 15 exceed even the 43% chimeric read estimate. A potential threshold for setting distance could be the mean bootstrap value of taxonomic identification. Note that the higher the minimum parent distance, the lower the mean bootstrap value of the removed ASVs. This indicates the threshold is doing a better job of selectively removing ASVs without a known Silva database resolved taxonomic ID; increasing the likelihood that the source of the removed ASV was technical.

**Table 5.**
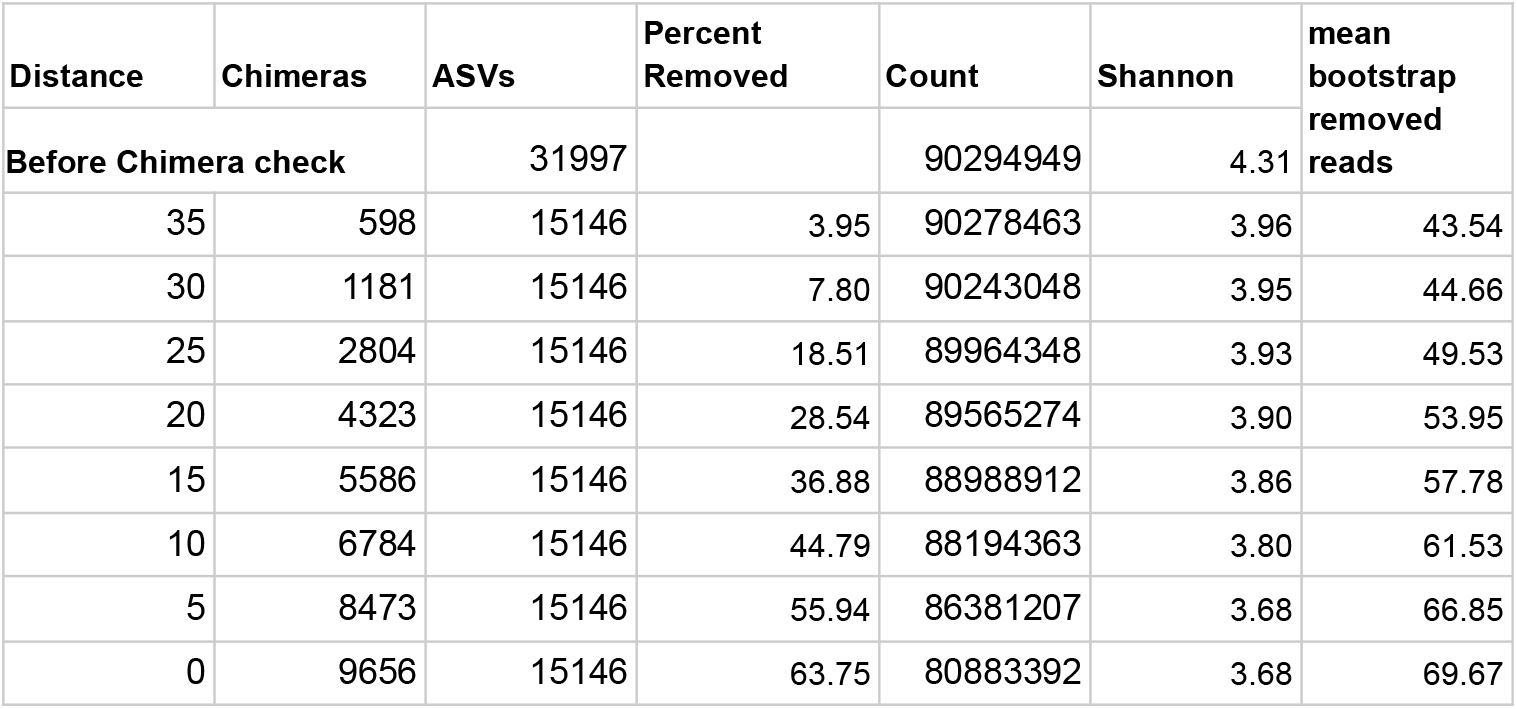
ASV dataset where a distance cutoff for potential parent ASVs is enforced for Bimeric sequence detection. As distance increases, the number of chimeras and percent removed is reduced. Average taxonomic identification confidence is lower as distance is increased.

### Applying Clustering

Applying the methods of clustering to Dada2 ASV tables before any filtering occurs reduces technical noise. Clustering Aviti_4 (Table 3), a human digestive study of 16S V4 region, with an initial unique ASV count of 10,839 to nearest 2 errors (99%) yields 5,986 ASVs total and synthetic mock ASVs decreased from 415 to 258. The mean count of ASVs in the synthetic mock sample goes from 56.8 to 102.87. Reducing the presence of PCR-induced ASVs makes it easier to isolate index swapped reads and contamination for a more accurate downstream analysis. Interestingly there are only 140 single sample ASVs remaining in this study after clustering reduced the initial 7,277 singletons. In comparison, clustering Aviti_1, a soil study of 16S V4 region with an initial unique ASV count of 246,750 to nearest 2 errors (99%) yields 78,931 ASVs total, a significant reduction in sample complexity. Similarly to the previous study, after clustering there are only 21 singleton ASVs remaining from an initial 115,321 singletons. All together this shows that clustering results in a significant noise reduction in the datasets that will help downstream analysis.

qPCR analysis of Aviti_1, the high singleton study samples, before PCR as compared to post sequencing statistics (Figure 4) shows an increased average number of singletons and counts in samples with more biological material before PCR; however sequence count and number of singletons have little to no relationship beyond samples with no qPCR signal have very low sequence count and those sequences that do exist for the sample are very likely to be singletons. qPCR methods are not a perfect measure of biological material and should be considered a rough estimate at best, especially at lower concentrations (Bustin et al). That said, the trend of more singletons beyond a minimum qPCR threshold of 1000 is clear; the question is whether that is an indication of biological signal or technical signal. Given that the singletons in traditionally higher signal material (soil and human) cluster together with higher count ASVs but remain in lower biomass material would point to a technical source.

**Figure 4.**
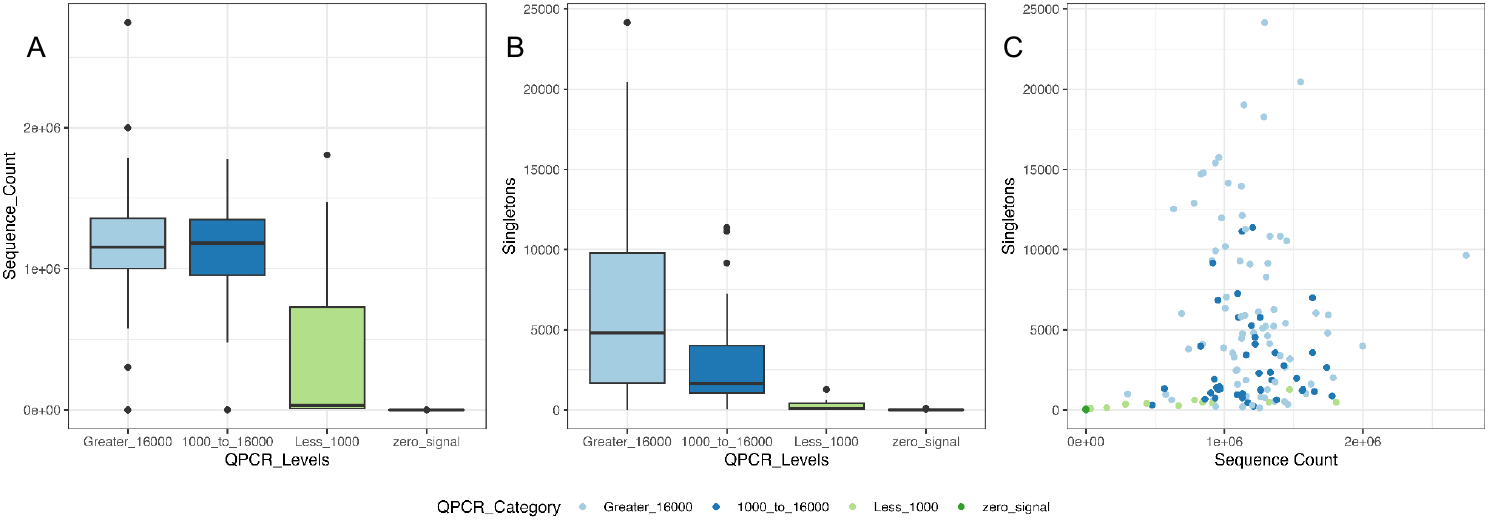
QPCR data. (A) Qpcr levels by sequence count show reduced count when samples <1000 sequence count. B) Higher QCPR concentrations are associated with increased singleton ASVs in samples. C) No relationship between Sequence count and singleton count.

Large increases of unique ASVs in Aviti data are a reflection of upstream steps in the sequencing process. The synthetic mock samples point specifically to errors introduced by the kapa hifi polymerase used in PCR (Shagin et al). Past work has pointed to DNA damage incurred in PCR cycling and other issues related to the PCR process (Potapov and Ong, McInerney et al.). Singleton ASV prevalence is related to the sample complexity and amount of starting biological material (Allen et al.). Further synthetic mock ASVs being introduced in greater numbers as a result of the length of the variable region isolated; again points to technical processes introducing non-biological variation.

## Conclusion

These analyses indicate that clustering Aviti projects in a count specific manner retains sequencing counts and corrects for some of the PCR induced errors. Limiting chimeric read detection based on parent distance is a necessary correction for Aviti read diversity and quality. The efficacy of these methods is dependent on the care used in the upstream pipeline and the amount and quality of the initial DNA material. The methods used upstream of sequencing, such as DNA extraction protocol and PCR, are going to be reflected in the final data more accurately with Aviti than previous technologies due to the massive upscaling in total reads per project. While some of this technical variation certainly existed in Illumina studies previously, it was likely hidden by the lower average quality of the sequencing data and lack of high-depth-per-sample-studies that are now available in Aviti datasets. Excitingly, Aviti’s high quality and large sequence count output allows researchers to ask questions not previously possible, but also increases the risk of the biological signal getting lost in the technical noise. This study provides a starting point for comparative analyses of Aviti marker gene sequencing data with those generated by other sequencing technologies.

## Data Availability

Marker gene datasets will be made public by contributing labs as research is completed. Code and synthetic mock datasets are available for analysis on: https://github.com/trevorjgould/Aviti_Processing

## Acknowledgments

Tim Heisel and The University of Minnesota Genomics Center for contribution of mock data and consultations.

Dr. Peter Kennedy (UMN) for suggestions on the manuscript.

Data Contributions from Hamilton lab, Santelli lab, Staley lab, Collister lab.

